# Conditional gene expression reveals stage-specific functions of the unfolded protein response in the *Ustilago maydis*/maize pathosystem

**DOI:** 10.1101/775692

**Authors:** Lara Schmitz, James W. Kronstad, Kai Heimel

## Abstract

*Ustilago maydis* is a model organism to study biotrophic plant-pathogen interactions. Sexual and pathogenic development of the fungus are tightly connected since fusion of compatible haploid sporidia is prerequisite for infection of the host plant, maize (*Zea mays*). After plant penetration, the unfolded protein response (UPR) is activated and required for biotrophic growth. The UPR is continuously active throughout all stages of pathogenic development *in planta*. However, since development of UPR deletion mutants stops directly after plant penetration, the role of an active UPR at later stages of development has/could not be examined, yet. Here, we establish a gene expression system for *U. maydis* that uses endogenous, conditionally active promoters to either induce or repress expression of a gene of interest during different stages of plant infection. Integration of the expression constructs into the native genomic locus and removal of resistance cassettes were required to obtain a wild type-like expression pattern. This indicates that genomic localization and chromatin structure are important for correct promoter activity and gene expression. By conditional expression of the central UPR regulator, Cib1, in *U. maydis*, we show that a functional UPR is required for continuous plant defense suppression after host infection and that *U. maydis* relies on a robust control system to prevent deleterious UPR hyperactivation.

## Introduction

The phytopathogenic basidiomycete *Ustilago maydis* causes the smut disease on maize (*Zea mays*) and is a well-established model organism to study sexual fungal development and biotrophic fungal/plant interactions, but also basic cellular processes such as DNA recombination and vesicular transport (Bakkeren *et al*., 2008; Banuett, 1995; Dean *et al*., 2012; Kahmann and Kämper, 2004; Lanver *et al*., 2018).

The available genome sequence, a broad range of molecular techniques and tools, as well as a highly efficient homologous recombination system enable the precise genetic manipulation of *U. maydis* (Brachmann *et al*., 2004; Kämper, 2004; Kämper *et al*., 2006; Schuster *et al*., 2016; Terfrüchte *et al*., 2014). Common and frequently used ways to characterize gene functions are available including deletion or overexpression of genes, as well as the generation of gene fusions for fluorescence microscopy or epitope tagging. PCR-based methods for gene replacement via homologous recombination as well as promoters for constitutive, inducible or titratable (over)expression of genes like the *tef*, *otef, nar1, crg1* or *tet-Off* promoter are also available (Banks *et al*., 1993; Bottin *et al*., 2002; Brachmann *et al*., 2004; Kämper, 2004; Spellig *et al*., 1996; Zarnack *et al*., 2008). These promoters can be fused to a gene of interest and are either integrated in the native gene locus or into the locus of the succinate dehydrogenase-encoding gene (*UMAG_00844, sdh2*; *ip* locus) by homologous recombination, conferring carboxin resistance (Keon *et al*., 1991). However, gene expression analysis using metabolism-dependent promoters may result in pleiotropic effects due to metabolic changes and unwanted overexpression of the gene of interest. Other conditional gene expression systems in fungi include for example estrogen-, orzearalenone-, or light-inducible expression systems for *Aspergillus sp.* (Pachlinger *et al*., 2005), *Gibberella zeae* (Lee *et al*., 2010), or *Neurospora crassa* (Salinas *et al*., 2018), respectively (see Kluge *et al*., 2018 for a comprehensive overview). These systems are all suitable to control gene expression under axenic culture conditions. However, tools to address the function of genes specifically during the process of organismal interactions, such as fungal/plant interactions, are not well established, yet.

*U. maydis* is a dimorphic fungus, specifically infecting its host plant maize. Sexual and pathogenic development are interconnected because plant infection requires cell/cell fusion of compatible haploid sporidia to generate the infectious, dikaryotic filament. Development of the fungus including mating, filamentous growth, plant penetration and biotrophic growth *in planta* are controlled by a tetrapolar mating-type system (Hartmann *et al*., 1996; Bölker, 2001; Feldbrügge *et al*., 2004; Wahl *et al*., 2010). The *a*-mating type locus encodes a pheromone-receptor system that regulates cell-cell recognition and fusion (Bölker *et al*., 1992), whereas all subsequent steps of pathogenic development are controlled by the bE/bW-heterodimer encoded by the *b*-mating type locus (Schulz *et al*., 1990; Kämper *et al*., 1995; Heimel *et al*., 2010a; Wahl *et al*., 2010). After penetration of the plant surface, *U. maydis* establishes a compatible biotrophic interaction with the host plant by secreting effectors that suppress plant defense reactions (Lanver *et al*., 2017; Lo Presti *et al*., 2015a). Expression of effector-encoding genes is specifically induced during the fungal/plant interaction (Kämper *et al*., 2006; Lanver *et al*., 2018), resulting in increased stress imposed on the endoplasmic reticulum (ER). Activation of the unfolded protein response (UPR) is critical to counteract elevated ER stress levels and for efficient secretion of effector proteins (Hampel *et al*., 2016; Pinter *et al*., 2019; Lo Presti *et al*., 2015b). The UPR is controlled by a key regulatory bZIP transcription factor termed Hac1 in *Saccharomyces cerevisiae*, XBP1 in higher eukaryotes and Cib1 in *U. maydis* (Cox and Walter, 1996; Heimel *et al*., 2013; Kawahara *et al*., 1998; Rüegsegger *et al*., 2001). The UPR is activated by unconventional cytoplasmic splicing of the *HAC1*/*cib1*/*XBP1* mRNA, generating the processed form of the mRNA (e.g. *cib1*^s^) that is translated into the active transcription factor. Hence, the effects of genetic UPR activation can be analyzed by expression of the *cib1*^s^ mRNA without drug induced side-effects.

In fungal human and plant pathogens, a functional UPR is necessary for disease development (Cheon *et al*., 2011; Heimel *et al*., 2013; Joubert *et al*., 2011; Kong *et al*., 2015; Richie *et al*., 2009; Yi *et al*., 2009). In *U. maydis*, the UPR is specifically activated after plant penetration and remains constantly active during all subsequent stages of biotrophic growth inside the host plant (Heimel *et al*., 2013). This suggests that the UPR is constantly required for efficient protein secretion and regulation of pathogenic growth. However, since *cib1* mutant strains are arrested early after plant infection, the relevance of a functional UPR at later stages of biotrophic development *in planta* could not be addressed, yet.

Here, we established a system for conditional and stage-specific gene expression during pathogenic growth of *U. maydis in planta*. Based on previously published time-resolved transcriptome data of fungal gene expression during biotrophic growth (Lanver *et al*., 2018), genes with desired *in planta* expression patterns were identified and their promoters were used for conditional gene expression. Importantly, we observed that maintenance of the genomic context and removal of resistance marker cassettes are required for correct promoter activity and conditional gene expression. To address the function of the UPR regulator Cib1 at later stages of biotrophic development, we used conditional promoters to repress, induce or overexpress *cib1* at specific stages of biotrophic growth *in planta*. We thereby demonstrate that *U. maydis* is resistant to UPR hyperactivation after plant penetration, suggesting effective strategies to prevent or cope with deleterious ER stress. By contrast, repression of *cib1* expression at 2 or 4 days post inoculation (dpi) revealed that a functional UPR is not only essential for establishment of biotrophy, but also required for colonization and continuous suppression of the plant defense at later stages of development *in planta*.

## Results

### Genomic localization and the presence of resistance marker cassettes affect the activity of promoters specifically expressed *in planta*

In previous studies, promoters of *U. maydis mig* (maize induced genes)-genes that are specifically expressed *in planta* were used for conditional gene expression during infection (Lo Presti *et al*., 2015b; Scherer *et al*., 2006; Wahl *et al*., 2010). In addition to the *mig1* gene (Basse *et al*., 2000), *mig* genes include the *mig2* gene cluster harboring five highly homologous genes, all of which are plant-specifically expressed but not involved in the virulence of *U. maydis* (Basse *et al*., 2002). The *mig2*-genes (*mig2_1, mig2_2, mig2_3, mig2_4 and mig2_5*) differ in their strength and temporal dynamics of expression. Thus, their promoters represent suitable targets for controlled and plant-specific expression/overexpression of a gene of interest.

To address the effect of overexpressing the spliced version of the *cib1* mRNA (*cib1^s^* in the following text), encoding the UPR regulator Cib1, during pathogenic development *in planta*, we integrated a P*_mig2_1_:cib1^s^* promoter fusion into the *ip* locus of the solopathogenic SG200 strain (Kämper *et al*., 2006). The *ip* or *cbx* locus is commonly used for integration of linear DNA into the *U. maydis* genome by homologous recombination, conferring resistance against carboxin (Brachmann, 2001). Since the virulence of strain SG200P*_mig2_1_:cib1^s^* was severely attenuated in plant infection experiments (Figure 1A), we investigated at which stage pathogenic development was blocked. Our analysis revealed the inability of SG200P*_mig2_1_:cib1^s^* to induce filamentous growth on charcoal containing solid media and on the leaf surface (Figure 1B), suggesting that pathogenic development is abrogated before plant penetration.

**Figure 1.**
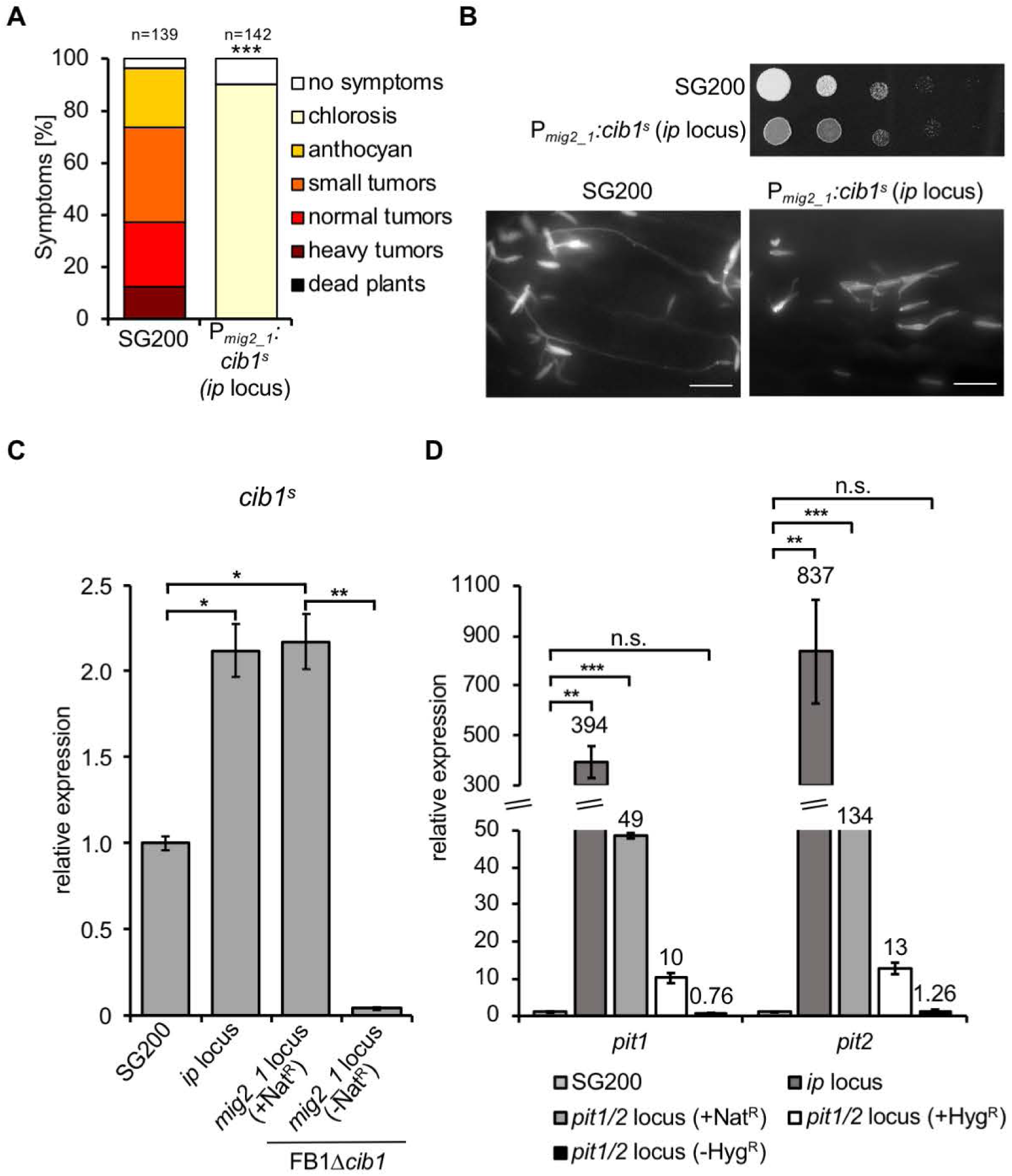
The locus of integration and presence of a resistance cassette influence promoter activity. **(A)** Plant infection assay with the solopathogenic strain SG200 and a derivative expression strain. Strains SG200 and SG200 P*_mig2_1_:cib1^s^* (*ip* locus) were inoculated into 8 day-old maize seedlings. Disease symptoms were rated 8 dpi and grouped into categories as shown in the figure legend. n = number of inoculated plants. Significance was calculated using the Mann-Whitney-test. ***P < 0.001 **(B)** Analysis of *b*-dependent filament formation on PD-CC solid media and on the leaf surface. Strains SG200 and SG200P*_mig2_1_:cib1^s^* (*ip* locus) were spotted on PD-CC solid media. Photographs were taken after 24 hours at 28°C. White fuzzy colonies indicate the formation of filaments. Fungal hyphae were stained 24 hours after inoculation with calcofluor to visualize the cells. Scale bar =10 µm. **(C)** qRT-PCR analysis of *cib1^s^* gene expression when integrated in different loci and after removal of the resistance cassette. Primers specifically detecting the spliced *cib1* transcript were used. RNA was isolated from exponentially growing *U. maydis* strains SG200, SG200 P*_mig2_1_:cib1^s^* (*ip* locus integration), FB1Δ*cib1*Δ*mig2_1*::P*cib1*^s^ (*mig2_1* locus, +Nat^R^) and FB1Δ*cib1*Δ*mig2_1*::*cib1*^s^ (*mig2_1* locus, -Nat^R^). *eIF2b* was used for normalization. Expression values represent the mean of three biological replicates with two technical duplicates each. Error bars represent the SEM. Statistical significance was calculated using the students *t* test. *P value < 0.05, **P < 0.01, and ***P < 0.001. **(D)** qRT-PCR analysis of *pit1* and *pit2* gene expression when integrated in different loci and after removal of the resistance cassette. RNA was isolated from exponentially growing *U. maydis* strains SG200, SG200 P*_pit1/2_:pit2/1* (*ip* locus integration), SG200 P*_pit1/2_:pit2/1* (*pit2/1* locus, +Nat^R^), SG200 P*_pit1/2_:pit2/1* (*pit2/1* locus, +Hyg^R^) and P*_pit1/2_:pit2/1* (*pit2/1* locus, -Hyg^R^). *eIF2b* was used for normalization. Expression values represent the mean of three biological replicates with two technical duplicates each. Error bars represent the SEM. Statistical significance was calculated using the students *t* test. *P value < 0.05, **P < 0.01, and ***P < 0.001.

We have previously shown that constitutive expression of *cib1*^s^ inhibits the formation of infectious filaments (Heimel *et al*., 2013). Hence, we tested if integration of the P*_mig2:1_*:*cib1*^s^ construct into the *ip* locus might result in increased expression levels of *cib1*^s^ during growth in axenic culture. Indeed, levels of *cib1*^s^ were significantly increased in strain SG200P*_mig2_1_:cib1^s^*, when compared to the SG200 control strain (Figure 1C). Since elevated *cib1*^s^ levels might either result from increased activity of the *cib1* wild type (WT) ORF that is also present in SG200P*_mig2_1_:cib1^s^*, or from “leaky” P*_mig2_1_-*driven expression, we used the Δ*cib1* background for further analyses. To study if this effect is specific for the *ip* locus, we generated *U. maydis* strain FB1Δ*cib1* Δ*mig2_1*::*cib1*^s^ (*mig2_1* locus (+Nat^R^)) by replacing the *mig2_1* ORF with the *cib1*^s^ gene. To exclude potential effects of the resistance cassette used for integration, the nourseothricin (Nat^R^) resistance cassette was removed by FLP/FRT recombination (Khrunyk *et al*., 2010). This revealed that elevated *cib1*^s^-levels indeed resulted from aberrant P*_mig2_1_* activity and only strains in which the nourseothricin resistance marker was removed (*mig2_1* locus (-Nat^R^)) were devoid of any detectable *cib1*^s^ expression (Figure 1C). In summary, our data strongly suggest that both the genomic locus and the presence of a resistance marker contribute to the increased activity of the *mig2_1* promoter in axenic culture.

To pinpoint if this effect is specific for *cib1^s^*, we performed an analogous experiment with the *pit1* and *pit2* genes, which are divergently transcribed from the same promoter. In axenic culture, expression of both genes is barely detectable but highly induced during biotrophic growth *in planta* (Doehlemann *et al*., 2011; Lanver *et al*., 2018). We determined expression levels of both genes when (re-)integrated into *U. maydis* strain SG200Δ*pit1/2* (Hampel *et al*., 2016) into 1) the *ip* locus or the native *pit1/2* locus, using either 2) nourseothricin (Nat^R^) or 3) hygromycin resistance (Hyg^R^) cassettes and 4) after removal of the resistance marker (Figure 1D). Surprisingly, transcript levels of both *pit1* and *pit2* were drastically increased when integrated into the *ip* locus (approximately 400-fold and 800-fold, respectively) in comparison to the SG200 (WT) control. Even when expressed from their native genomic locus, transcript levels of both genes were still significantly increased (*pit1*: 49-fold (Nat^R^) and 10-fold (Hyg^R^); *pit2*: 134-fold (Nat^R^) and 13-fold (Hyg^R^)) and only after removal of the resistance marker cassette (*pit1/2* locus (-Hyg^R^)) expression of *pit1* and *pit2* was similar to the SG200 (WT) control (Figure 1D). In summary, these data demonstrate that the locus of integration and the presence of resistance marker cassettes influence the activity of “conditional promoters”.

### Overexpression of *cib1^s^* does not disturb pathogenic development *in planta*

To set up a system that allows for proper functioning of conditional promoters we constructed plasmids harboring promoters of the *mig1*, *mig2_1*, *mig2_2* or *mig2_3* genes. 3’ sequences were followed by a *Sfi*I restriction site for integration of the gene of interest, an FRT-Hyg^R^ or an FRT-Nat^R^ resistance marker cassette and a 1kb sequence harboring the 3’ UTR for recombination and integration into the genomic locus of respective *mig* genes. It is important to note that neither single nor the combined deletion of all *mig* genes negatively affected pathogenic development of *U. maydis* (Farfsing *et al*., 2005). To specifically increase *cib1*^s^ levels *in planta* and address the effect of UPR hyperactivation on pathogenic development, we expressed *cib1*^s^ under control of the *mig1* or the *mig2_1* promoter. To this end, the *mig1* or *mig2_1* ORFs were replaced by *cib1^s^*, followed by the removal of the resistance marker cassette in the *U. maydis* strain FB1Δ*cib1* (Heimel *et al*., 2010b) (see Supplemental Figure 1 for an overview of the approach). We first checked for leaky *cib1*^s^-expression by testing ER stress resistance and filamentous growth of the generated strains. When spotted on solid media supplemented with the ER stress-inducing drugs tunicamycin (TM) or dithiothreitol (DTT), the hyper-susceptibility of the FB1Δ*cib1* progenitor strain was not suppressed, suggesting that P*_mig1_* and P*_mig2_1_* are not active in axenic culture (Figure 2A). Consistently, filamentous growth of respective strain combinations was not affected in mating assays on charcoal containing potato dextrose (PD) solid media (Figure 2B), thus confirming the absence of leaky *cib1*^s^ expression.

**Figure 2.**
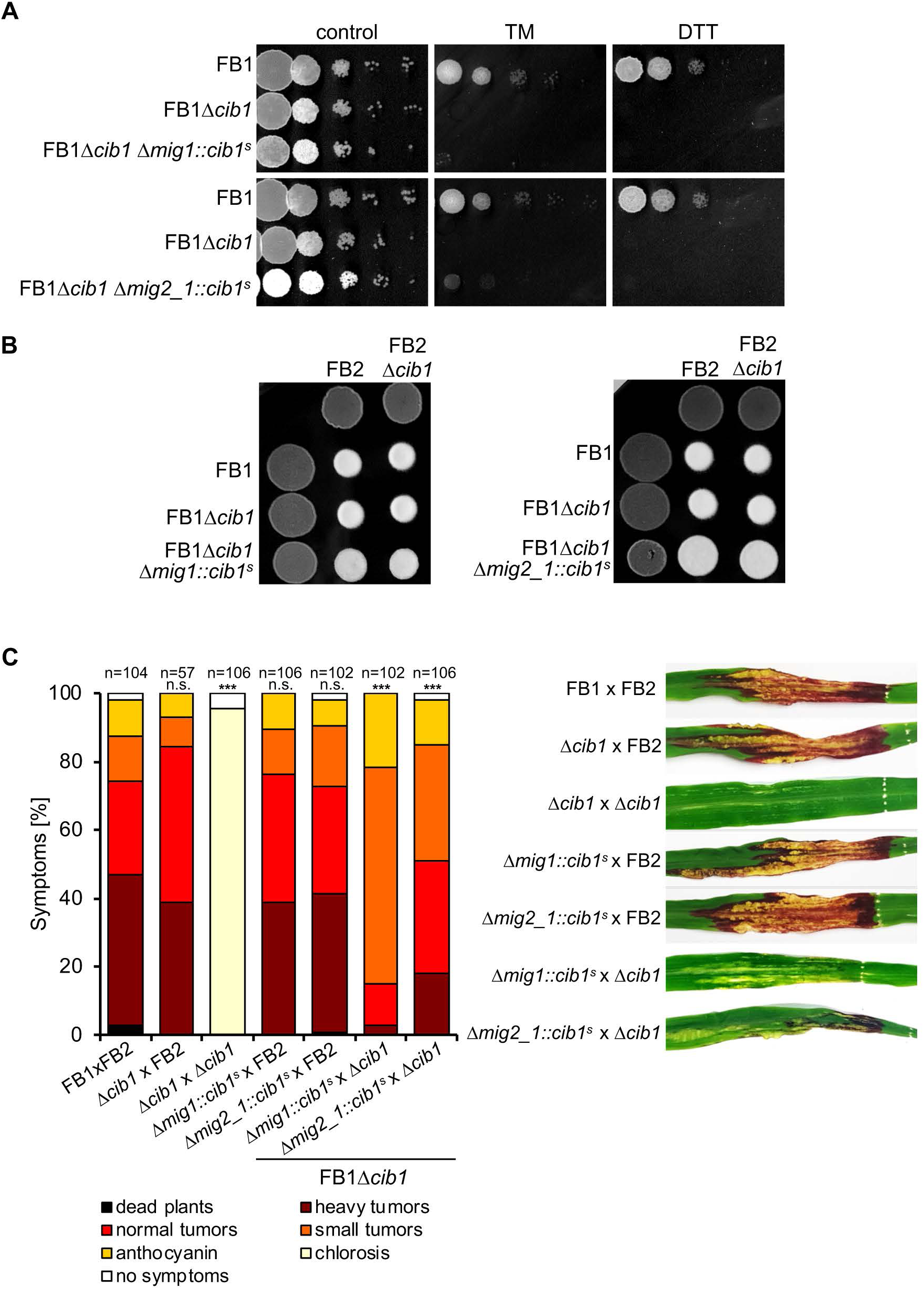
Overexpression of *cib1*^s^ *in planta* does not affect pathogenicity of *U. maydis*. **(A)** ER stress assay of strains FB1, FB1*Δcib1*, FB1*Δcib1* Δ*mig1*::*cib1*^s^ and FB1*Δcib1* Δ*mig2_1*::*cib1*^s^. Serial 10-fold dilutions were spotted on YNBG solid medium supplemented with TM (1.0 μg/ml) or DTT (1 mM). Pictures were taken after 48 hours of incubation at 28 °C. **(B)** Mating assay with compatible mixtures of FB1, FB2, FB1Δ*cib1*, FB2Δ*cib1*, FB1*Δcib1* Δ*mig1*::*cib1*^s^ and FB1*Δcib1* Δ*mig2_1*::*cib1*^s^. Mixtures were spotted on PD-CC solid media as shown in the figure. Photographs were taken after 24 hours at 28°C. White fuzzy colonies indicate the formation of filaments. **(C)** Plant infection assay with compatible mixtures of FB1 and FB2, FB1Δ*cib1*, FB2Δ*cib1*, FB1Δ*cib1*Δ*mig1*::*cib1*^s^ and FB1Δ*cib1* Δ*mig2_1*::*cib1*^s^. 8 day-old maize seedlings were co-inoculated with the indicated strain mixtures. Disease symptoms were rated 8 dpi and grouped into categories as shown in the figure legend. n = number of inoculated plants. Pictures of leaves were taken at 8 dpi and represent the most common infection symptom. Significance was calculated using the Mann-Whitney-test. ***P < 0.001.

Mixtures of mating compatible strains FB1, FB2, FB1Δ*cib1*, FB2Δ*cib1*, and the derivatives FB1Δ*cib1*Δ*mig1*::*cib1*^s^ and FB1Δ*cib1*Δ*mig2_1*::*cib1*^s^ were used for plant infection studies (Figure 2C). P*_mig1_*- or P*_mig2_1_*-mediated expression of *cib1*^s^ did not affect pathogenicity when strains were combined with the compatible FB2 WT strain. By contrast, when FB1Δ*cib1*Δ*mig1*::*cib1*^s^ or FB1Δ*cib1*Δ*mig2_1*::*cib1*^s^ were combined with the compatible FB2Δ*cib1* deletion mutant, virulence was strongly increased compared to the non-pathogenic FB1Δ*cib1* × FB2Δ*cib1* control, although not to WT (FB1 × FB2) levels. This result suggests that the mechanisms to prevent UPR hyperactivation *in planta* are robust and efficient in *U. maydis* thereby confirming the previous assumption that the UPR is specifically required during biotrophic development *in planta* (Heimel *et al*., 2010b; Heimel *et al*., 2013).

### Establishment of a system for *in planta*-specific gene depletion

We next aimed to establish a gene expression system that would allow us to examine gene functions during defined developmental stages *in planta* by using promoters that are specifically repressed during plant infection. To this end, we screened the publicly available RNAseq data set published by Lanver *et al*., 2018 and identified a total of four candidate genes which are expressed during axenic growth and early steps of pathogenic development before plant penetration, but strongly repressed shortly after plant penetration (1-2 dpi; *UMAG_00050*, *UMAG_05690* and *UMAG_12184*,), or at later stages during biotrophic growth *in planta* (*UMAG_03597*) (Lanver *et al*., 2018).

We focused on *UMAG_12184* and *UMAG_03597* for our current studies. Both genes are expressed in axenic culture and at early stages of pathogenic development, but are strongly repressed at 2 (*UMAG_12184*) or 4 dpi (*UMAG_03597*) (Figure 3A), during and shortly after *U. maydis* has established a compatible biotrophic interaction with its host plant. To test if these genes are involved in virulence, we deleted the genes in the haploid, solopathogenic *U. maydis* strain SG200. SG200 expresses a compatible bE1/bW2-heterodimer, and is thus capable of forming filaments and infecting its host plant, *Z. mays*, without the need of a compatible mating partner (Kämper *et al*., 2006). Both deletion strains were not affected in virulence (Figure 3B), demonstrating that these genes are dispensable for pathogenic development. In addition, neither ER or cell wall stress resistance, nor filamentous growth on charcoal containing PD solid media were strongly affected by either deletion, although filament formation was reduced in the *UMAG_03597* deletion mutant (Supplemental Figure 2). However, since SG200Δ*UMAG_03597* showed full virulence, this phenotype does not impair the ability of the fungus to cause disease. Based on these results, the respective promoters were regarded as suitable candidates to be used for conditional gene expression.

**Figure 3.**
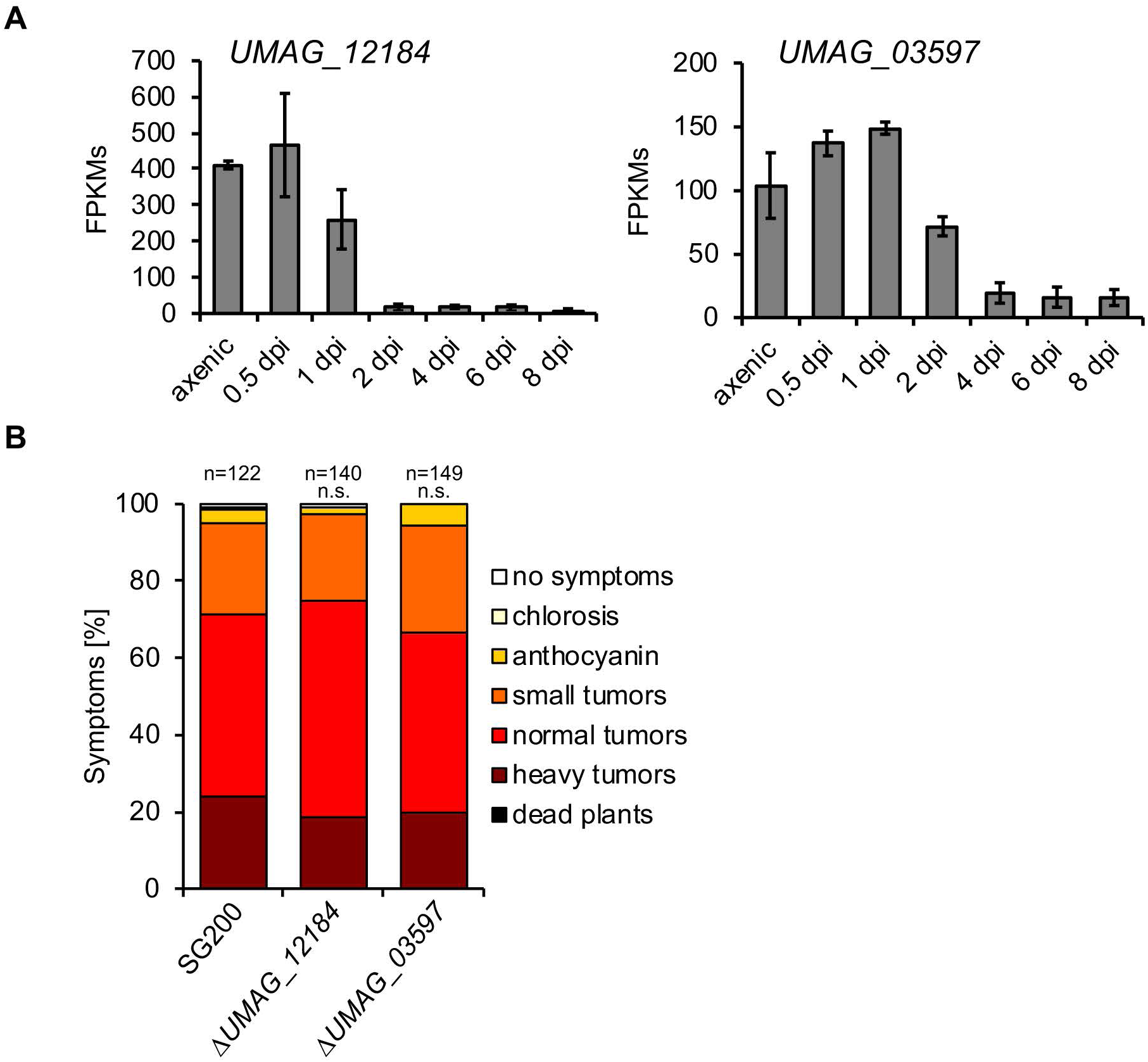
Identification and testing of promoters for conditional gene expression. **(A)** Fragments Per Kilobase Million (FPKMs) of the *UMAG_12184* and *UMAG_03597* genes up to 8 days post inoculation (dpi). 6 day-old maize seedlings were injected with a mixture of compatible haploid strains FB1 and FB2 and plant material was harvested at the indicated time points. Raw data was extracted from Lanver *et al*., 2018. **(B)** Plant infection assay with the solopathogenic strain SG200 and derivatives. SG200, SG200*ΔUMAG_12184* and SG200*ΔUMAG_03597* were inoculated into 8 day-old maize seedlings. Disease symptoms were rated 8 days after inoculation (dpi) and grouped into categories as shown in the figure legend. n = number of inoculated plants. Significance was calculated using the Mann-Whitney-test.

### Cib1 is required throughout biotrophic development *in planta*

The bZIP transcription factor Cib1 is the central regulator of the UPR in *U. maydis*, and required for coordinating pathogenic development, efficient secretion of effectors and plant defense suppression (Heimel *et al*., 2013; Pinter *et al*., 2019). Pathogenic development of *cib1* deletion strains is blocked immediately after plant penetration resulting in the complete absence of tumor formation (Heimel *et al*., 2013). To test if *cib1* is only important directly after plant penetration (e.g., for release of the cell cycle block and establishment of the biotrophic interaction), or if it is also necessary at later stages of pathogenic development, we expressed *cib1* under control of the *UMAG_12184* and *UMAG_03597* promoters (shut off at 2 and 4 dpi, respectively). To this end, we replaced *UMAG_12184* or *UMAG_03597* genes with the *cib1* ORF in strain FB2Δ*cib1* (Heimel *et al*., 2010b), generating strains FB2Δ*cib1* Δ*UMAG_12184::cib1* and FB2Δ*cib1* Δ*UMAG_03597::cib1*. Resistance cassettes used for selection of successful integration events were removed by FLP/FRT mediated recombination (Khrunyk *et al*., 2010).

The generated strains were tested for correct expression of *cib1* under axenic conditions by ER stress assays using TM or DTT. Both mutants showed ER stress resistance similar to the WT (FB2) control, demonstrating that *cib1* expression driven by either promoter is sufficient to suppress the ER-stress hypersensitivity of the FB2Δ*cib1* progenitor strain (Figure 4A) (Heimel *et al*., 2013). Additionally, when compatible mixtures of WT (FB1 × FB2), Δ*cib1* derivatives (FB1Δ*cib1* × FB2Δ*cib1*) or derivatives expressing *cib1* under control of conditional promoters (FB1Δ*cib1* × FB2Δ*cib1*Δ*UMAG_12184::cib1* or FB1Δ*cib1* × FB2Δ*cib1*Δ*UMAG_03597::cib1*) were spotted on charcoal containing PD solid media (Figure 4B), all tested combinations developed white fuzzy colonies (Banuett and Herskowitz, 1989) indicating that mating is not affected in these strains.

**Figure 4.**
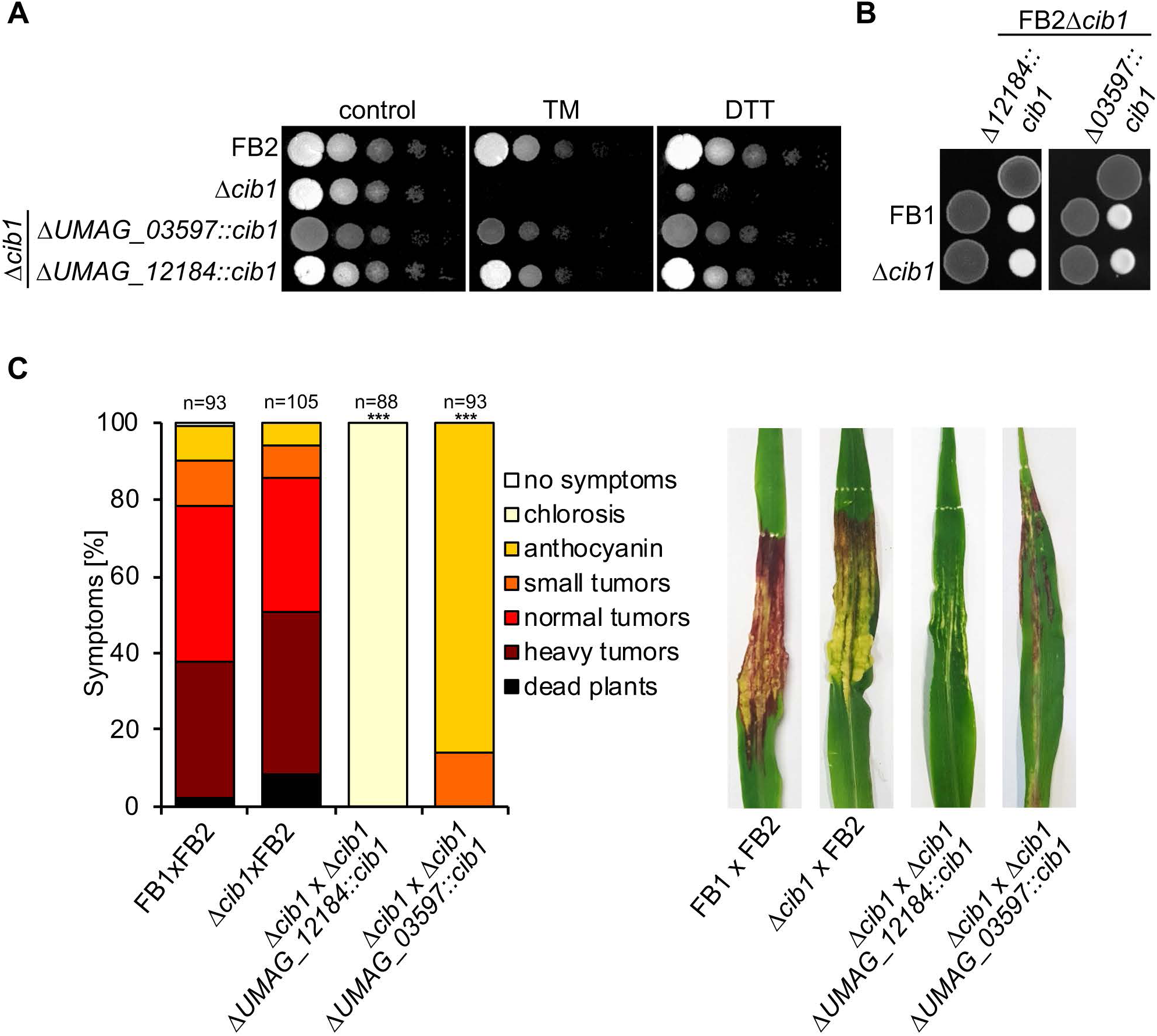
Conditional *cib1* expression restores ER-stress resistance, but not pathogenicity. **(A)** ER stress assay of strains FB2 (WT), FB2*Δcib1*, and derivatives. Serial 10-fold dilutions were spotted on YNBG solid medium supplemented with TM (1.0 μg/ml) or DTT (1.0 mM). Pictures were taken after 48 hours of incubation at 28 °C. **(B)** Mating assay with FB1, FB1Δ*cib1* and FB2Δ*cib1* Δ*UMAG_12184::cib1* and FB2Δ*cib1* Δ*UMAG_03597::cib1*. Compatible mixtures of strains were spotted on potato dextrose solid media supplemented with 1% charcoal (PD-CC). Photographs were taken after 24 hours at 28°C. White fuzzy colonies indicate the formation of filaments. **(C)** Plant infection assay with FB1 and FB2, FB1*Δcib1* and FB2, FB2Δ*cib1* Δ*UMAG_12184::cib1* and FB2Δ*cib1* Δ*UMAG_03597::cib1*. 8 day-old maize seedlings were co-inoculated with compatible strain mixtures as indicated in the figure. Disease symptoms were rated 8 dpi and grouped into categories as shown in the figure legend. n = number of inoculated plants. Pictures of leaves were taken at 8 dpi and represent the most common infection symptom. Significance was calculated using the Mann-Whitney-test. ***P < 0.001

Next, we investigated the effect of plant-specific repression of *cib1* in plant infection assays. When compatible mixtures of FB1Δ*cib1* × FB2 strains were used for inoculation of maize plants, symptom development was indistinguishable from the WT (FB1 × FB2) control (Figure 4C), demonstrating that a single functional copy of *cib1* is sufficient for full virulence of the fungus. However, when *cib1* was expressed under the control of P*_UMAG_12184_* (FB1Δ*cib1* × FB2Δ*cib1*Δ*UMAG_12184::cib1*), virulence was almost completely abolished and no tumors were formed, resembling the Δ*cib1* phenotype. By contrast, expression of *cib1* under the control of P*_UMAG_03597_* (FB1Δ*cib1* × FB2Δ*cib1*Δ*UMAG_03597::cib1*) was sufficient to trigger anthocyanin production and the formation of small tumors. This indicates that prolonged expression of *cib1* is sufficient to overcome the developmental block of Δ*cib1* strains, and initiate pathogenic growth *in planta*.

To visualize fungal growth *in planta* and assess at which step biotrophic development of the fungus stopped, infected leaves were harvested at 2, 4 and 6 dpi and stained with Chlorazol Black E (Figure 5A). Microscopic analysis revealed extensive proliferation and clamp cell formation when plants were inoculated with combinations of WT (FB1 × FB2) or FB1 × FB2Δ*cib1* strains. When *cib1* was expressed under the control of P*_UMAG_12184_* until 2 dpi (FB1Δ*cib1* × FB2Δ*cib1*Δ*UMAG_12184::cib1*) infectious dikaryotic filaments penetrated the plant surface via appressoria at 2 dpi, but did not progress further in the plant at later stages (4 and 6 dpi). Consequently, clamp cell formation and extended fungal proliferation was not observed. By contrast, expression of *cib1* under control of P*_UMAG_03597_:cib1* (FB2Δ*cib1*Δ*UMAG_03597::cib1*) enabled the fungus to overcome the cell cycle block and induce proliferation, as reflected by hyphal branching and the formation of clamp cells at 4 dpi. However, the subsequent colonization of host tissue by fungal hyphae at 6 dpi appeared strongly reduced in comparison to the controls (FB1 × FB2 and FB1Δ*cib1* × FB2) (Figure 5A). This suggests that the reduced activity of P*_UMAG_03597_* and the resulting decrease of *cib1* levels at this stage prevents further progression of fungal hyphae inside the plant.

**Figure 5.**
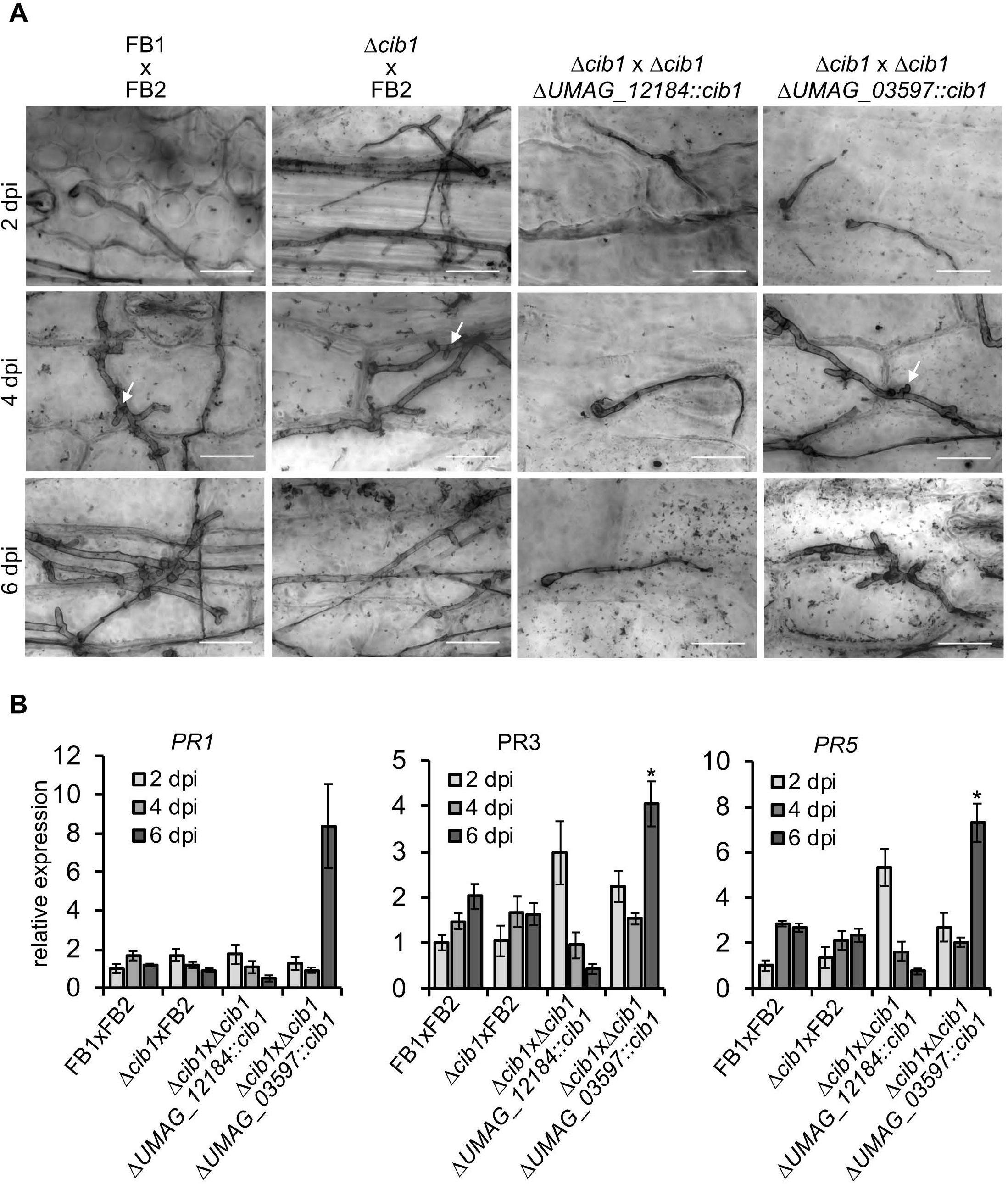
Analysis of fungal morphology and plant defense response of conditional *cib1* mutant strains. **(A)** Fungal proliferation of compatible mixtures of FB1 and FB2, FB1Δ*cib1* and FB2, FB2Δ*cib1* Δ*UMAG_12184::cib1*or FB2Δ*cib1* Δ*UMAG_03597::cib1*investigated by Chlorazol Black E staining of infected leaf samples at 2, 4 and 6 dpi. Arrows point to clamp cells indicative of fungal proliferation *in planta*. Scale bar = 20 µm. **(B)** qRT-PCR analysis of *PR1, PR3* and *PR5* gene expression of infected maize leaves at 2, 4 and 6 dpi. Maize seedlings were inoculated with the indicated strains. *GAPDH* was used for normalization. Expression values represent the mean of two or three biological replicates with two technical duplicates each. Error bars represent the SEM. Statistical significance was calculated using the students *t* test. *P value < 0.05.

Previous studies revealed that plants inoculated with Δ*cib1* mutant strains show increased plant defense reactions as demonstrated by elevated expression of pathogenesis related (*PR*) gene expression at 2 dpi (Heimel *et al*., 2013). It is conceivable that this observation is connected to the requirement of a functional UPR for efficient secretion and processing of effectors (Lo Presti *et al*., 2015b; Hampel *et al*., 2016; Pinter *et al*., 2019). To investigate if Cib1 is also required for plant defense suppression at later stages, we determined expression levels of *PR* genes *PR1*, *PR3 and PR5* at 2, 4 and 6 dpi in plants inoculated with strains conditionally expressing *cib1*. All three *PR* genes are markers for salicylic acid (SA)-related defense responses that are typically suppressed by biotrophic plant pathogens like *U. maydis* (Glazebrook, 2005). Consistent with the results obtained in infection studies, P*_UMAG_12184_*-driven expression of *cib1* resulted in increased expression of *PR3 and PR5* genes at 2 dpi, whereas expression of *PR1* was not induced (Figure 5B). By contrast, when *cib1* was expressed under the control of P*_UMAG_03597_*, expression of all three *PR* genes was induced at 6 dpi. These observations are consistent with the expected activity of the P*_UMAG_12184_* and P*_UMAG_03597_* promoters that are repressed at 2 and 4 dpi, respectively. Hence, our data indicate that *cib1* expression under control of the promoter of *UMAG_12184* is not sufficient to establish a compatible biotrophic interaction *in planta* leading to a block in pathogenic development. By contrast, when *cib1* is expressed for an extended time (from promoter P*_UMAG_03597_*), a compatible interaction appears to be established, allowing further proliferation. This suggests that *cib1* is required for plant defense suppression not only at the onset (2 dpi), but also during later (4 and 6 dpi) stages of biotrophic development *in planta*.

## Discussion

Analysis of gene function typically involves the generation of gene deletion and overexpression strains. To test for functions related to the virulence of plant pathogenic fungi, deletion strains are inoculated into the host plant and scored for development of disease symptoms (Dean *et al*., 2012). However, the analysis of virulence factors that are essential for pathogenic development relies on the description of the first phenotype that is observed, i.e. the stage when pathogenic development is blocked. Hence, potential functions of these factors that might also be important at later stages of pathogenic development have not been addressed and remain elusive. To date, suitable tools to address this problem are restricted to the introduction of a gate keeper mutation in kinases that can be chemically inhibited by non-hydrolyzable ATP analogs. However, this strategy is only suitable for the analysis of kinase functions and requires extensive controls to exclude potential side-effects of the chemical treatment (Sakulkoo *et al*., 2018).

In this study, we report a conditional gene expression system for *U. maydis* that enables the study of gene functions at different stages of pathogenic development in the plant. We identified suitable promoters that are active during axenic growth and repressed during pathogenic growth *in planta*. We demonstrate that promoters (e.g., P*_mig2_1_* or P*_pit1/2_*), previously used for plant-specific gene expression, are active during axenic growth and produce considerable amounts of transcripts (up to 800-fold induced expression for *pit2*) when integrated into the *ip* locus or when resistance marker cassettes are located in their vicinity. Proper promoter function required the maintenance of the genomic environment by “in locus” integration (as demonstrated for the *mig2_1* or *pit1/2* genes) and removal of the resistance marker cassette.

Similar to the *mig2* gene cluster, the virulence factors *pit1* and *pit2* are part of a gene cluster that is specifically upregulated *in planta* (Basse *et al*., 2002; Doehlemann *et al*., 2011). Interestingly, gene expression of the majority of effector gene clusters including *mig2*-and *pit*-clusters is induced in strains deleted for the histone deacetylase *hda1* (Reichmann *et al*., 2002; Treutlein, 2007), suggesting that these clusters are subject to epigenetic regulation. It remains to be investigated if this effect is restricted to clustered effector genes or accounts for the regulation of non-clustered effectors as well. Chromatin-based regulation of effector genes appears to be a common feature in plant pathogenic fungi (Soyer *et al*., 2014). It is well established that the RNA polymerase II complex closely interacts with histone modifying enzymes, including the SWI/SNF complex and histone acetyltransferases (Wittschieben *et al*., 1999; Wittschieben *et al*., 2000). This complex is supposed to function as a chromatin snowplow leading to increased accessibility of the genomic neighborhood (Barton and Crowe, 2001). Hence, although the underlying molecular details remain to be addressed, it is tempting to speculate that high expression of the *sdh2* gene (*ip* locus), or of highly expressed resistance marker genes might affect the chromatin structure and thus de-repress silent promoters in their vicinity.

The conditional overexpression of *cib1*^s^ using the *mig1* or *mig2_1* promoter did not result in alterations of disease symptoms. Because the *mig1* promoter is highly active *in planta* (Basse *et al*., 2000; Lanver *et al*., 2018), it is especially remarkable that high levels of *cib1*^s^ are not detrimental for fungal proliferation *in planta*. This suggests that *U. maydis* has established effective control mechanisms to prevent UPR hyperactivation, one of which is based on the functional modification of the UPR by the Cib1/Clp1 interaction, providing ER stress hyper-resistance of Clp1-expressing strains (Heimel *et al*., 2013; Pinter *et al*., 2019). A potential second mechanism might be reminiscent of UPR regulation in higher eukaryotes and involve the unspliced *cib1* transcript, or the encoded Cib1^u^ protein (Heimel *et al*., 2013). In higher eukaryotes, the U-isoform of the Hac1-like UPR regulator XBP1 functions as a repressor of the UPR (Yoshida *et al*., 2006). Hence, a similar mode of action would potentially counteract increased *cib1*^s^ levels, as expression of the unspliced *cib1* transcript itself is subject to Cib1-dependent gene regulation.

The increasing body of transcriptomic data provides a highly valuable treasure box to identify promoters with desired expression dynamics. In theory, this enables establishment of tailor-made expression systems to address gene specific functions in a sophisticated manner. However, our attempt to identify promoters that are active during axenic growth, but strongly repressed at different stages of pathogenic development *in planta* revealed only a low number of candidates. Moreover, we observed that it is desirable for correct promoter function to maintain the genomic context. Using Cib1, an essential virulence factor in *U. maydis*, we carried out a proof-of-principle analysis demonstrating that a functional UPR is not only required directly after penetration of the leaf surface (Heimel *et al*., 2010b; Heimel *et al*., 2013), but also at later stages of pathogenic development. The increased expression of *PR* genes correlates with repression of promoter activity and thus reduced *cib1* transcript levels. This strongly suggests that continuous suppression of the SA-related plant defense depends on sustained UPR activity. This is consistent with the observation that not only early but also late effectors require the UPR for efficient secretion and/or processing (Lo Presti *et al*., 2015b; Hampel *et al*., 2016; Pinter *et al*., 2019). Although our system is applicable for a wide range of genes, a potential limitation is met when examining stage-specific functions of genes with dynamic expression patterns. One way to enable these studies would be the stage specific expression of site-specific recombinases, such as CRE or FLP (Sadowski, 1995; Sternberg and Hamilton, 1981; Sauer and Henderson, 1988), as established for a variety of model systems including numerous fungi (Khrunyk *et al*., 2010; Kopke *et al*., 2010; Kück and Hoff, 2010; Mizutani *et al*., 2012; Twaruschek *et al*., 2018; Zhang *et al*., 2013). In this way, loxP or FRT flanked genes could be targeted for genomic deletion in a stage- or development-specific manner, while maintaining their dynamic expression pattern.

In summary, we established a conditional expression system that allows one to address plant-specific functions of genes of interest in the *U. maydis*/maize pathosystem. The generation of constructs to be integrated into the genome is facilitated by an efficient one step cloning procedure. Plasmids for conditional induction or repression of genes during biotrophic development *in planta* are cross compatible and harbor identical *Sfi*I restriction sites for easy exchange of genes. Since the constructs can either be integrated into the genome of solopathogenic or compatible haploid strains, future studies using combinations of conditionally expressed constructs will allow the consideration of even more sophisticated scientific questions, such as the relevance of posttranslational modifications or enzymatic activity of a protein for biotrophic growth of *U. maydis*.

## Experimental Procedures

### Strains and Growth Conditions

*Escherichia coli* TOP10 strain was used for cloning and amplification of plasmid DNA. *U. maydis* cells were grown at 28°C in YEPS light medium (Tsukuda *et al*., 1988), complete medium (CM) (Holliday, 1974) or yeast nitrogen base (YNB) medium (Freitag *et al*., 2011; Mahlert *et al*., 2006). Mating assays were performed as described before (Brachmann *et al*., 2001). ER-stress assays were carried out on YNB solid media containing the indicated concentrations of DTT or TM (Sigma-Aldrich). Sensitivity to Calcofluor White or Congo red was tested by drop-assay on YNB solid media containing the indicated concentration of the respective compound. Filamentous growth assays were carried out using potato-dextrose (PD) media supplemented with 1% charcoal (PD-CC) (Holliday, 1974). Strains used in this study are listed in Supplemental Table 1.

### DNA and RNA procedures

Molecular methods followed described protocols (Sambrook *et al*., 1989). For gene deletions, a PCR-based approach was used (Kämper, 2004). Isolation of genomic DNA from *U. maydis* and transformation procedures were performed according to Schulz *et al*., 1990. Homologous integration was performed using linearized plasmid DNA or PCR-amplified DNA. Integration was verified by Southern hybridization. Total RNA was extracted from exponentially growing cells in axenic culture using Trizol reagent according to the manufacturer’s instructions (Invitrogen, Karlsruhe, Germany). RNA integrity was checked by agarose-gel-electrophoresis. Residual DNA was removed from total RNA samples using the TURBO DNA-*free*™ Kit (Ambion, Darmstadt, Germany). cDNA was synthesized using the iScript™ cDNA Synthesis Kit (BioRad, Munich, Germany). Primers used in this study are listed in Supplemental Table 2.

### Quantitative RT-PCR

qRT-PCR analysis was performed as described (Hampel *et al*., 2016). For all qRT-PCR experiments, three independent biological replicates and two technical replicates were analyzed using the MESA GREEN qPCR MasterMix plus for SYBR Assay with fluorescein (Eurogentech, Cologne, Germany). qRT-PCR was performed using the CFX Connect Real-Time PCR Detection System and analyzed with the CFX Manager Maestro Software (BioRad).

### Plasmid construction

For gene deletions, a PCR-based approach and the *SfiI* insertion cassette system were used (Brachmann *et al*., 2004; Kämper, 2004). For construction of plasmids for conditional gene expression, 0.5-1 kb flanking regions of chosen genes (*UMAG_03597, UMAG_12184, mig1, mig2_1*) were PCR amplified from genomic DNA, adding a *SfiI* restriction site to the 5’ of the left border (LB) and a *BamH*I (for *UMAG_12184, mig1* and *mig2_1*) or *Kpn*I (for *UMAG_03597*) restriction site to the 3’end of the right border (RB). The gene of interest (GOI; *cib1* or *cib1^s^*) was PCR amplified from genomic DNA or from plasmid P*_cib1_:cib1^s^*, respectively, adding *SfiI* restriction sites to the 5’ and 3’end. The Hyg^R^ cassette was amplified from plasmid pUMa1442 adding a *BamH*I (for *UMAG_12184*) or *Kpn*I restriction site (for *UMAG_03597*) to the 3’ end and a *Sfi*I restriction site to the 5’ end. The resulting DNA fragments were ligated to obtain *LB-GOI-Hyg^R^-RB* or *LB-GOI-Nat^R^-RB* and integrated into the pCR2.1 TOPO vector (Invitrogen) or the pJet1.2 vector (ThermoFisher Scientific, Waltham, Massachusetts, USA) according to the manufacturer’s instructions to generate plasmids pCR2.1 P*_UMAG_12184_*:*cib1*(NatR), pCR2.1 P*_UMAG_03597_*:*cib1*(HygR), pJet1.2 P*_mig2_1_*:*cib1^s^*(NatR) and pJet1.2 P*_mig1_*:*cib1^s^*(NatR).

For construction of the P*_mig2_1_:cib1^s^* construct for *ip* locus integration, the vectors pMig2_1:clp1 and pRU11-cib1s were cut with *NdeI* and *EcoRI*. The resulting 2.0 kb *cib1^s^* fragment of pRU11-cib1s (Heimel *et al*., 2013) and the 5.2 kb backbone of Mig2_1:clp1 were ligated to obtain plasmid P*_mig2_1_:cib1^s^*. Plasmids generated in this study are listed in Supplemental Table 3.

### Plant Infections

The haploid, solopathogenic strain SG200 and its derivatives or FB1 and FB2 and their respective derivatives were grown to an OD600 of 0.6-0.8 in YEPS light medium, adjusted to an OD600 of 1.0 in water and mixed 1:1 with a compatible mating partner. The resulting suspension was used to inoculate 8-day-old maize seedlings of the variety Early Golden Bantam. Plants were grown in a CLF Plant Climatics GroBank (Wertingen, Germany) with a 14 h (28°C) day and 10 h (22 °C) night cycle. Symptoms were scored according to disease rating criteria reported by Kämper *et al*., 2006. Three independent clones were used for each plant infection experiment and the average scores for each symptom are shown in the respective diagrams. Photographs from infected leaves were taken and represent the most common infection symptoms for the respective mutant.

### Chlorazole Black E staining and microscopy

Infected leaf tissue was harvested at 2, 4 and 6 dpi and kept in 100% ethanol until further processing. Chlorazole Black E staining was performed as described in Brachmann *et al*., 2001. Microscopic analysis was performed using an Axio Imager.M2 equipped with an AxioCam MRm camera (ZEISS, Jena, Germany). All images were processed using ImageJ.

### Quantification of *U. maydis* gene expression *in planta* and *PR* gene expression

Infected leaf tissue was harvested at the indicated time points. Samples of five infected maize seedlings were pooled per replicate, frozen in liquid nitrogen and ground to powder by mortar and pestle according to Lanver *et al*., 2018. Total RNA was extracted using Trizol reagent (Invitrogen) and used for qRT-PCR analysis as described above. For expression analysis of *U. maydis* genes, *eIF2b* expression levels were used for normalization. Expression of *PR1*, *PR3* and *PR5* from *Zea mays* were determined and normalized to *GAPDH* expression levels.

### Statistical Analysis

Statistical significance was calculated using Student’s *t* test. The statistical significance of plant infection phenotypes was calculated using the Mann-Whitney test as described previously (Freitag *et al*., 2011). Results were considered significant if the P value was <0.05.

### Accession numbers

Sequence data from this article can be found in the National Center for Biotechnology Information database under the following accession numbers: *UMAG_12184*, XP_011388913.1; *UMAG_03597*, XP_011390022.1; *cib1, UMAG_11782*, XP_011390112.1; *mig2_1, UMAG_06178*, XP_011392548.1; *mig1, UMAG_03223*, XP_011389652.1; *pit1, UMAG_01374*, XP_011387263.1, *pit2, UMAG_01375*, XP_011387264.1; *PR1* (Zm.15280.1), BM351351; *PR3* (Zm.1085.1), BM339391; *PR5* (Zm.6659.1), BM075306; *GAPDH* (NM001111943).

## Acknowledgements

We thank Daniel Lanver and Regine Kahmann for sharing expression data prior to publication and Ivo Feussner for generous support. All authors acknowledge funding by the Deutsche Forschungsgemeinschaft (DFG), and the Natural Sciences Engineering Council of Canada (NSERC) through the International Research Training Group 2172 PRoTECT and a Discovery Grant (to JWK).

## Data availability statement

The data that support the findings of this study are available from the corresponding author upon reasonable request.

## Supporting Information Legends

**Supplemental Figure 1: Strategy for strain generation for conditional gene expression.**

1) The gene of interest (GOI) is deleted from its native genomic locus. 2) The GOI is integrated into the genomic locus of the conditionally expressed gene, thereby replacing the native gene. 3) The resistance marker (here: Nat^R^) is removed using the FLP/FRT recombination system.

**Supplemental Figure 2:** Δ*UMAG_12184* and Δ*UMAG_03597* strains do not show increased sensitivity to cell wall- or ER-stresses.

Cell wall and ER-stress assays, and tests for filamentous growth of strains SG200, SG200Δ*cib1*, SG200Δ*UMAG_12184* and SG200Δ*UMAG_03597*. Serial 10-fold dilutions were spotted on YNBG solid media supplemented with Congo Red (100 µg/ml) or Calcofluor White (50 µM) to induce cell wall stress, and on YNBG solid media supplemented with TM (1.0 μg/ml) or DTT (1 mM) to induce ER-stress. Cells were spotted on PD-CC solid media to induce filamentous growth. Pictures were taken after 48 hours of incubation at 28 °C.

**Supplemental Table 1: Strains used in this study**

**Supplemental Table 2: Primers used in this study**

**Supplemental Table 3: Plasmid used in this study**

