## Supplementary figures and images for "Conditional gene expression reveals stage-specific functions of the unfolded protein response in the *Ustilago maydis*/maize pathosystem"

### Supplemental Figure 1

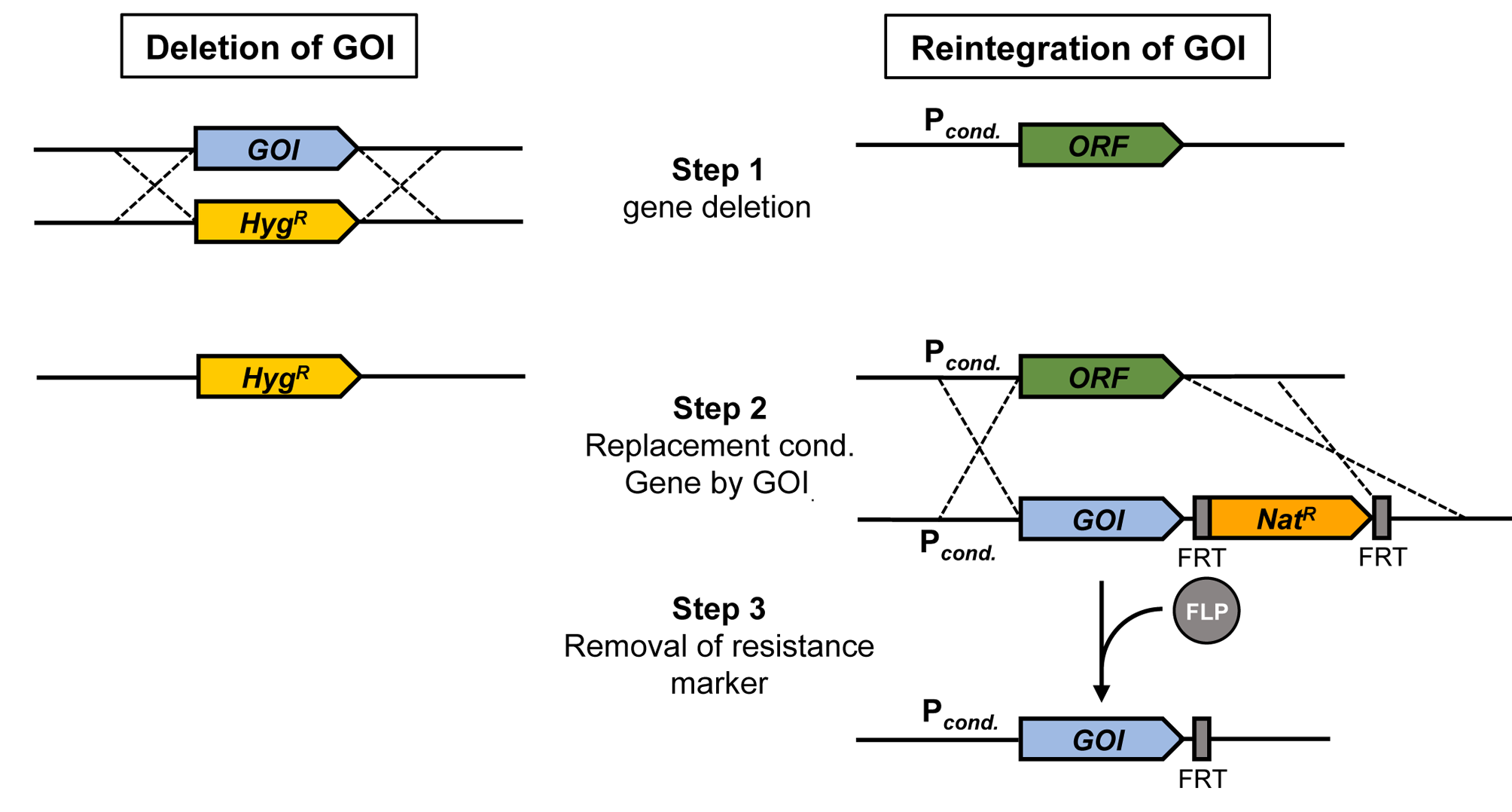

### Supplemental Figure 2

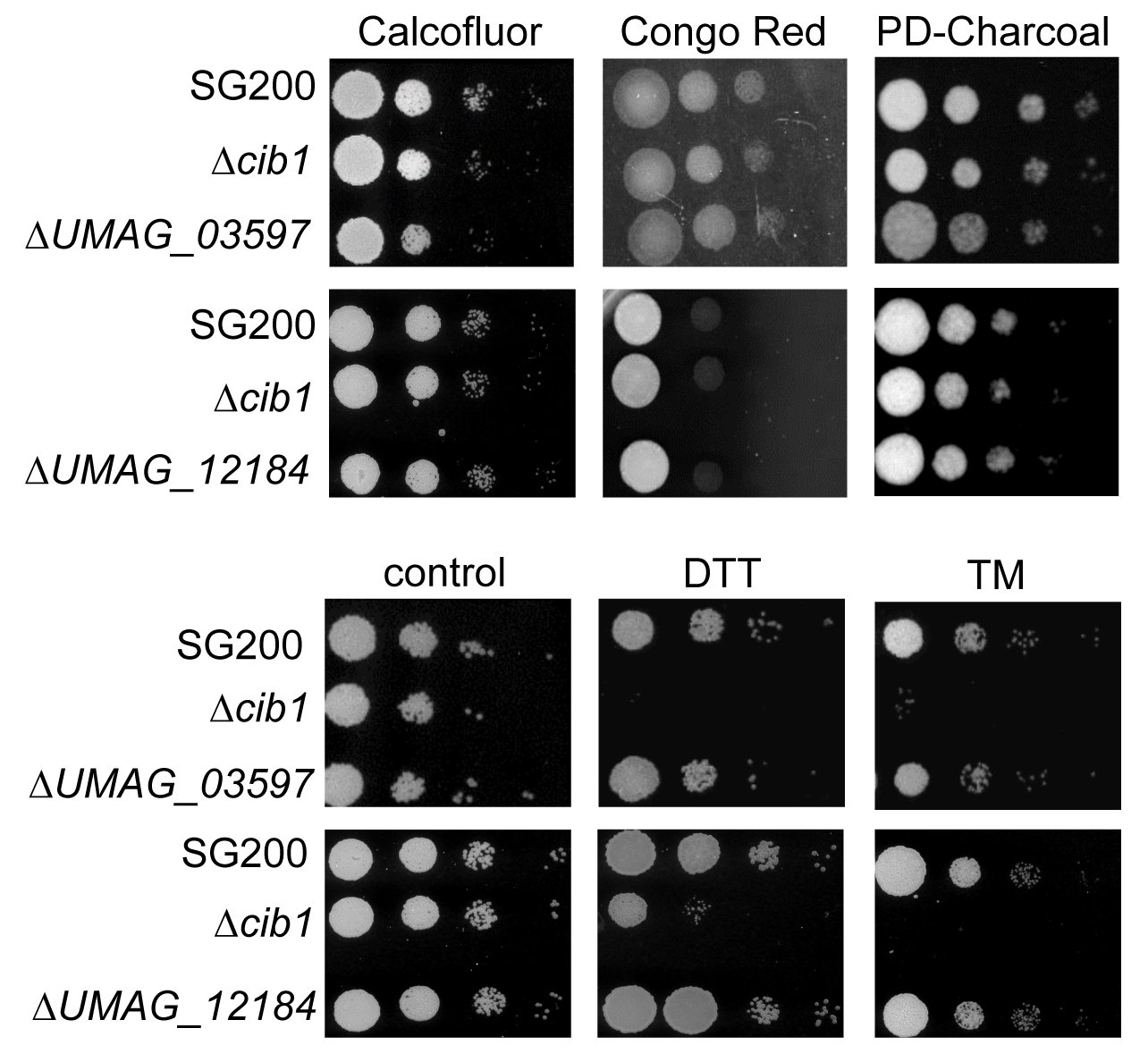
